# A novel sorting signal for RNA packaging into small extracellular vesicles

**DOI:** 10.1101/2023.06.08.544286

**Authors:** Yuma Oka, Kosei Tanaka, Yuki Kawasaki

## Abstract

Extracellular vesicles (EVs) play a critical role in the transport of functional RNAs to target recipient cells in numerous physiological processes. The RNA profiles present in EVs differed significantly from those in the originating cells, suggesting selective and active loading of specific RNAs into EVs. Small EVs (sEVs) obtained by stepwise ultracentrifugation have been reported to contain non-sEV components. Analysis of sEVs separated from non-sEVs components revealed that microRNAs may not be released by sEVs. This has raised interest in other RNA types, such as mRNA, which may be functional molecules released by sEVs. However, the molecular mechanisms underlying selective loading of mRNA into sEVs remain unclear. Here, we show that the part of 3’ untranslated region (UTR) sequence of *RAB13* selectively enriches RNA in sEVs and serves as an RNA signal for loading into sEVs. Our results demonstrate that *RAB13* is the most enriched RNA in sEVs, and this enrichment is primarily driven by its 3’UTR sequence. These findings highlight the potential of the *RAB13* 3’UTR sequence as an RNA signal that enables the loading of target RNA into sEVs. This technology has the potential to improve EV-based drug delivery and other applications.

## Introduction

Extracellular vesicles (EVs), which are released from diverse cell types, are critical for intercellular communication and play essential roles in many physiological processes^1^. EVs transfer various molecular cargoes, including RNAs, proteins, and metabolites that perform specific functions in the recipient cells^2,3^. The RNA profiles in EVs often differ significantly from those in their corresponding cells of origin^4^, suggesting an active and selective mechanism for RNA sorting into EVs.

Small EVs (sEVs) constitute a subclass of EVs that includes the types of vesicles classically known as exosomes, which are derived from endosome, are 50-150 nm in diameter, and are enriched with tetraspanin proteins such as CD9 and CD63. Stepwise ultracentrifugation is often performed to purify sEVs, and ultracentrifugal fractions contain other non-sEVs components called nonvesicular (NV) compartments^5^. High-resolution density gradient fractionation of sEVs and NVs has shown that microRNAs (miRNAs) are predominantly associated with the NV fractions rather than with the purified sEVs fractions, and proteins related to miRNA biogenesis, such as Argonaute 1-4, are absent from the sEVs fractions, suggesting that extracellular miRNAs may not be released by sEVs. Thus, other types of RNA, such as mRNA among cargo RNAs are of greater interest in terms of the functional molecules released by sEVs. However, the molecular mechanisms by which selective mRNAs are loaded into sEVs have not been fully elucidated. Some RNA motif sequences have been proposed to facilitate loading into sEVs; however, not all mRNAs enriched in sEVs possess these sequences.

The objective of this study was to elucidate the RNA signal sequences responsible for their sorting and packaging into sEVs. We employed a previously established immunoaffinity purification method to isolate CD9 positive EVs^6^. Transcriptome analysis revealed the enrichment of multiple mRNA species within EVs. Subsequent investigations identified the RNA region responsible for selective mRNA enrichment within EVs.

## Results

To obtain ultrapure sEVs from HEK293 cells, crude sEV, which is denoted as P100, the pellet resulting from centrifugation at 100,000 × g, were further fractionated into low-density sEVs and high-density non-vesicular components (NV) using sucrose density gradient ultracentrifugation (Supplementary Fig. S1 online). The fractions were separated into two groups, low- and high-density fractions, based on the particle concentrations determined by nanoparticle-tracking analysis (Supplementary Fig. S1 online). The low-density fractions (1.10-1.15 g/mL) showed the presence of sEVs markers CD9 and CD63, signifying that they contained sEVs (Fig. 1a). In contrast, the high-density fraction did not contain CD9 or CD63. The average particle size of each fraction was less than 150 nm, which is within the typical size range of sEVs (Supplementary Fig. S1 online). These results suggested that the particles present in the low-density fractions were CD9 positive sEVs. EV preparation by ultracentrifugation may cause aggregation of vesicles^7,8^. To obtain CD9 positive EVs without ultracentrifugation, CD9 positive EVs were directly isolated from the cell-conditioned medium using a previously established immunoaffinity purification method. The presence of CD9 and CD63 in purified CD9 positive EVs, as well as in P100, was confirmed (Fig. 1b).

**Fig. 1.**
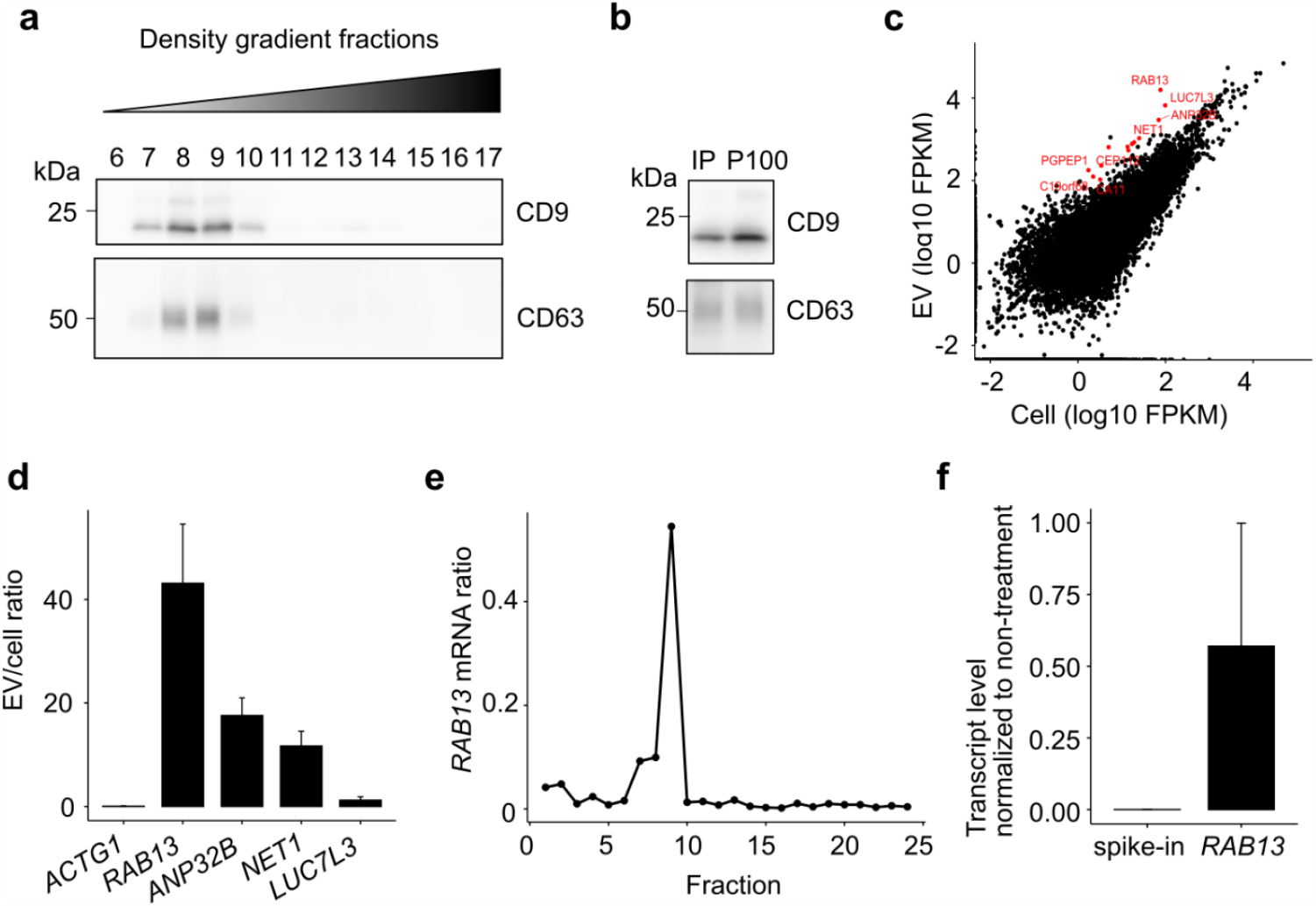
*RAB13* is enriched in sEVs. (a) Western blot of sucrose density gradient fractions extracted from conditioned medium of HEK293 cells. (b) Western blot of CD9 positive EVs purified by immunoaffinity purification. (c) Comparison of transcript levels between cells and CD9 positive EVs. The genes enriched in the CD9 positive EVs are shown in red. (d) Transcript enrichment level in CD9 positive EVs measured by RT-qPCR. The transcripts of interest in the CD9 positive EVs and cells were quantified and normalized to *GAPDH*, expressed as a ratio of transcript level in the CD9 positive EVs to that in the cells. (e) *RAB13* mRNA level in sucrose density gradient fractions was measured by RT-qPCR. The transcript level of *RAB13* was measured in each fraction and normalized to the total transcript level in all fractions. (f) Transcript levels for RNase-treated *RAB13* and spike-in control in P100 were measured by RT-qPCR. The transcript levels were normalized to non-treated samples.

RNA sequencing analysis of poly(A) RNA extracted from the CD9 positive EVs identified 13 transcripts with Fragments Per Kilobase of transcript per Million mapped reads (FPKM) over 100 in the CD9 positive EVs, 2^5^-fold higher than in the cells (Fig. 1c). Quantitative polymerase chain reaction (qPCR) was performed to validate the enrichment of specific transcripts in CD9 positive EVs. The ratio of RNA levels of *RAB13, ANP32B*, and *NET1* in EV to those in cells was over 10, indicating that they were highly enriched in CD9 positive EVs. In contrast, the enrichment ratio of *LUC7L3* was 1.2, comparable to that of the housekeeping gene *ACTG1* (Fig. 1d).

In the subsequent analyses, we focused on the most enriched RNA, *RAB13*. We investigated the localization of *RAB13* in sEVs in sucrose density gradient fractions and found that 73% *RAB13* was collected from fractions 7 to 9, with fraction 9 containing 54% (Fig. 1e). To further investigate whether *RAB13* was present inside the sEVs, we added spike-in RNA to P100 and performed RNase treatment to quantify the amount of RNA protected from RNase inside the membrane-bound vesicles. Our results demonstrated that 57% of *RAB13* RNA was present in P100 after RNase treatment, whereas less than 0.1% of the spike-in RNA was present in P100 after RNase treatment (Fig. 1f). These results indicated that *RAB13* RNA was indeed present inside the sEVs.

To determine the region within *RAB13* sequence necessary for sorting RNA into CD9 positive EVs, we established cell lines expressing partial sequences of *RAB13. RAB13* transcript was 1164 nucleotide long, with a coding sequence spanning positions 104–715 (Fig. 2a). Enrichment level in CD9 positive EVs showed that only the 3’-untranslated region (UTR) -containing transcripts (375-1164 and 783-1164) were enriched in the CD9 positive EVs (Fig. 2b). The sequence from positions 575 to 1164, which included the 3’UTR, was enriched by approximately 2-fold in CD9 positive EVs, whereas the transcripts without the 3’UTR region showed less than 2-fold enrichment in CD9 positive EVs.

**Fig. 2.**
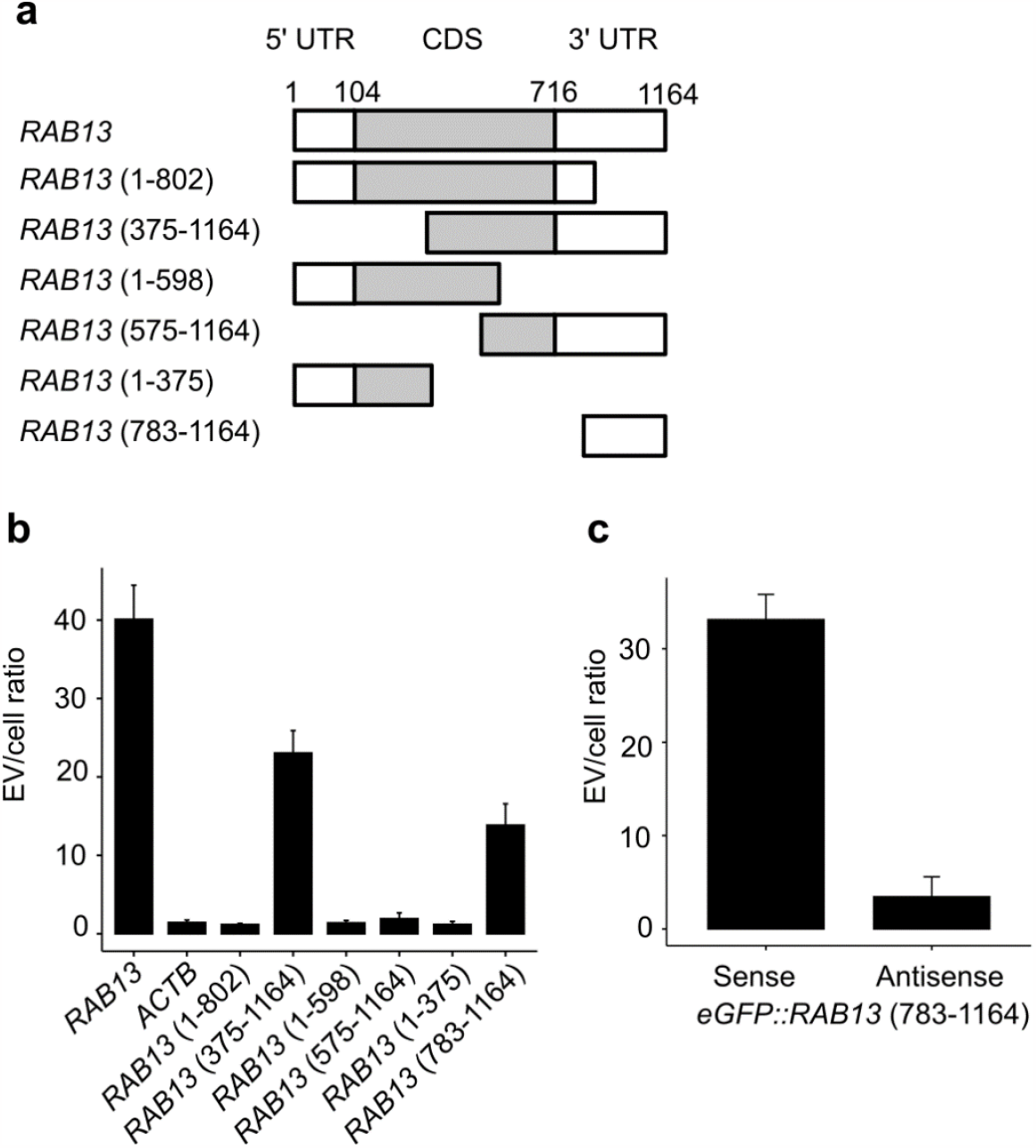
RNA loading into CD9 positive EVs occurs due to part of *RAB13* 3’UTR sequence. (a) Diagram of the *RAB13* mRNA and its partial sequences. The indicated number in parentheses corresponds to the sequence positions of *RAB13*. (b) Enrichment level of transgenes in the CD9 positive EVs from cells expressing *ACTB* or partial sequences of *RAB13*. (c) Enrichment level of transgenes fused with *RAB13* 3’UTR. The sequence of *RAB13* from position 783 to 1164 were fused with the 3’ end of *eGFP* and expressed in HEK293 cells.

To investigate the possibility of artificially sorting a transgene fused with *RAB13* 3’UTR sequence into CD9 positive EVs, we generated a transgene comprising of enhanced green fluorescent protein *(eGFP)* fused to the *RAB13* (783-1164) sequence in its 3’ end, in HEK293 cells. Analysis of the transcripts expressed in these cells and in the CD9 positive EVs showed 15-fold enrichment of the fusion gene in CD9 positive EVs (Fig. 2c). As a control, we also generated a fusion gene in which the 3’ end of *eGFP* was fused with the antisense strand of the *RAB13* (783-1164) sequence. This transcript was not enrichment in CD9 positive EVs.

We investigated the association between RNA stability and EV enrichment levels in endogenous RNAs globally. To evaluate RNA stability, direct RNA sequencing was performed to measure the poly(A) tail length of the poly(A) RNA extracted from cells expressing the *eGFP* fusion gene containing the *RAB13* (783-1164) sequence. The results showed no correlation between the poly(A) tail length and CD9 positive EVs enrichment for endogenous RNAs (r = −0.08330; Fig. 3a). Moreover, the median poly(A) tail length of *eGFP::RAB13* (783-1164) transcripts was 144.1 (Fig. 3b).

**Fig. 3.**
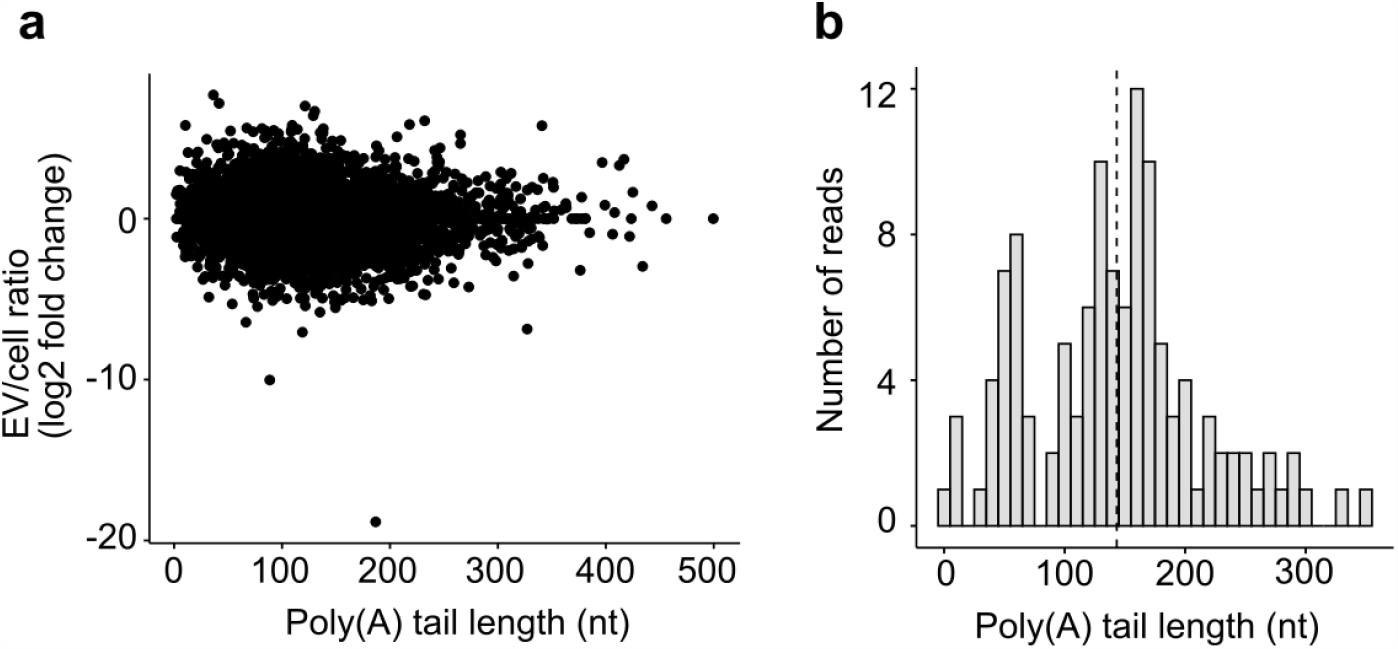
Association of poly(A) tail length with EV enrichment level. (a) Scatter plot showing the correlation between endogenous genes’ poly(A) tail length and their enrichment level within the CD9 positive EVs. (b) Poly(A) tail length distribution for the transgene extracted from cells expressing *eGFP::RAB13* (783-1164).

## Discussion

In the present study, we investigated the cargo RNA of CD9 positive EVs to identify the RNA sequences responsible for their loading into these EVs. We used an established immunoaffinity purification method to highly purify CD9 positive EVs from HEK293 cells. Transcriptome analysis revealed a high enrichment of RNA such as *RAB13, ANP32B*, and *NET1* in EVs. Notably, we found that the part of 3’UTR of *RAB13* acted as a loading signal for CD9 positive EVs, with an EV enrichment ratio of over 10, while non-enriched RNA, such as *ACTG1*, had an EV/cell ratio of around 1. The presence of non-enriched RNA in EVs may be attributed to passive loading or contamination with cellular debris that cannot be removed during EVs purification.

We found that *RAB13* was highly enriched in low-density sEVs isolated using sucrose density gradient fractionation as well as in CD9 positive EVs obtained through immunoprecipitation with anti-CD9 antibodies. Our results indicated the presence of heterogeneity within the sEVs population obtained through sucrose density gradient fractionation. Specifically, 73% of *RAB13* mRNA was preferentially localized in low-density fractions and 54% of *RAB13* mRNA was detected in fraction 9. This implies that a distinct subpopulations of sEVs exist and that each subpopulation may have unique characteristics that dictate its cargo. Importantly, our study highlights the critical role of EVs purification methods, such as high-resolution density gradient fractionation and immunoaffinity purification, in understanding the cargo of sEVs. To further investigate the biological functions and underlying mechanisms of sEVs, it is crucial to develop reliable methods for isolating and purifying specific sEV subpopulations. Establishing such methods would greatly enhance our understanding of the biological diversity of sEVs and their potential clinical applications.

Our study demonstrated the enrichment of *RAB13* in CD9 positive EVs from multiple cell lines, including HEK293, H1277, and MCF7 (Supplementary Fig. S2 online). Similar enrichment of *RAB13* has been reported in EVs derived from osteoblasts and colorectal cancer cell lines, albeit without a focus on purified sEVs^9,10^. This indicated that the mechanism responsible for the observed enrichment of *RAB13* may not be cell type-specific. Indeed, the low tissue specificity of *RAB13* expression may contribute to its enrichment in the EVs of various cell types.

It has been proposed that RNA transported to EVs possesses specific sequence motifs^11^. We analyzed the sequence to investigate whether any of the five reported RNA motif sequences (ACCAGCCT, CAGTGAGC, TAATCCCA, ACCCTGCCGCCTGGACTCCGCCTGT, and C[TA]G[GC][AGT]G[CT]C[AT]GG[GA]) were present in *RAB13*. However, none of these motifs were found in the *RAB13* sequence, suggesting that *RAB13* transport is not mediated by a recognition system that utilizes these motifs. It has been suggested that the motif-based loading system is cell type-specific^4^, and the cell lines employed in this study may possess a mechanism that recognizes and transports different sequence motifs. Furthermore, we were unable to detect any other human gene sequences homologous to the *RAB13* 3’UTR sequence, indicating its specificity to *RAB13*. Therefore, it is conceivable that *RAB13* and other enriched RNA could be loaded into EVs via distinct recognition mechanisms. The EV enrichment level did not show any correlation with the poly(A) tail length of endogenous gene transcripts from cells expressing either the sense or scrambled sequences of *RAB13* (783-1164) within the *eGFP* fusion gene (Supplementary Fig. S3 online). The median poly(A) tail length for the scrambled sequence was 78.0 with a read number of 37, which was insufficient for a comprehensive comparative analysis (Supplementary Fig. S3 online). The poly(A) tail of the fusion gene was not particularly long among the endogenous genes, although it was highly enriched in sEVs. Despite a low read count of 15, the median poly(A) tail length of the endogenous *RAB13* transcripts was recorded as 37.3. This does not constitute an exceptionally long poly(A) tail length among the endogenous genes. Collectively, these findings suggest that the poly(A) tail length is not a ubiquitous determining factor in the sorting of endogenous RNAs into sEVs, or specifically for *RAB13*. Our study revealed that the truncated 3’ end *RAB13* 3’UTR sequence (783-1037) showed similar enrichment level to the untruncated 3’ end *RAB13* 3’UTR sequence (783-1164), whereas the shorter subsequences of the *RAB13* 3’UTR (783-953 or 869-1037) were not enriched in sEVs by more than 2-fold (Supplementary Fig. S4 online), suggesting that sequences 783-868 and/or 954-1037 were vital for RNA loading. Further studies aimed at identifying molecules involved in transport are necessary to fully understand the mechanism of *RAB13* loading into sEVs.

Combining multiple regions of *RAB13* sequences may enhance loading efficiency, although this approach was not tested in our study. Our observations revealed that full-length sequences and longer subsequences (375-1164) exhibited higher enrichment in CD9 positive EVs compared to the 3’UTR subsequence (783-1163), while the intermediate sequence (575-1164) showed lower enrichment efficiency. These findings suggest that combining multiple sequences, such as the 5’ UTR sequence and the 3’UTR sequence, may potentially increase the loading efficiency.

Our study demonstrated that the fusion gene of the *RAB13* 3’UTR subsequence can be transported to sEVs. The *RAB13* 3’UTR subsequence has the potential to serve as an RNA loading technology for sEVs. Although several methods have been reported for transporting RNAs to EVs^12–14^, the use of *RAB13* sequences could complement these approaches. Currently, we do not observe eGFP fluorescence in recipient cells when HEK293 cells stably expressing an *eGFP* gene fused with the *RAB13* 3’UTR subsequent are co-cultured with wild-type HEK293 cells. Therefore, we could not confirm whether the RNA loaded into the sEVs migrated to and functioned in the recipient cells, at least when the recipient cells were HEK293 cells. However, endogenous *RAB1*3 and other enriched genes, *ANP32B* and *NET1*, were present in sEVs, indicating that these RNA may retain their function as transcripts even after transfer to sEVs and subsequent release into recipient cells (Supplementary Fig. S5 online). Further studies are required to investigate the behavior of RNA-loaded sEVs in different recipient cell types.

In conclusion, our study demonstrates that the *RAB13* 3’UTR subsequence can specifically enrich RNA in sEVs. The identified RNA region can be used in RNA-loading technologies, contributing to drug discovery and other applications.

## Methods

### Cell line and culture

Human embryonic kidney 293 cells (HEK293, #CRL-1573, ATCC) were maintained in Minimum Essential Medium alpha (MEMα) supplemented with 10% fetal bovine serum (FBS) and penicillin-streptomycin. The cells were cultured in a 5% CO_2_ humidified incubator at 37°C.

### Small EVs purification using ultracentrifugation

The conditioned medium from the cells was centrifuged at 2,000 × g for 5 min at 4°C, and then centrifuged at 10,000 × g for 15 min at 4°C. The obtained supernatant was layered onto 30 w/v% sucrose solution in ultracentrifugation tubes and subjected to ultracentrifugation at 100,000 × g for 3 h at 4°C using SW32Ti rotor (Beckman courter). The collected sucrose solutions were mixed with PBS and washed by ultracentrifugation at 100,000 × g for 1 h. The resulting pellets were used as crude sEVs and referred to as P100 in this study.

### Preparation of sucrose density gradient ultracentrifugation and fractionation

To prepare the sucrose gradient solutions, 1.4 mL of 2M sucrose in 1X PBS, 1 mL of 1.5 M sucrose in 1X PBS, and 1.4 mL of 1M sucrose in 1X PBS were layered into an ultracentrifugation tube. A 1 mL sample of crude EVs collected by ultracentrifugation was layered on top of the sucrose gradient solutions. The solutions were ultracentrifuged at 150,000 × g for 20 h at 4°C using P55ST2 rotor (Himac). Twenty-four fractions of 200 μL were collected from the top of the gradient. To determine the fraction densities, identical gradients with 1X PBS instead of the samples were generated using the same method. The fraction densities were measured using a PAL-BX/RI refractometer (Atago). Each collected fraction was mixed with 1X PBS and ultracentrifuged at 100,000 × g for 1 h. The pellets were resuspended in PBS.

### Immunoaffinity purification of CD9 positive EVs

The culture medium was mixed with in-house anti-CD9 antibody-conjugated magnetic beads and EV-binding buffer (0.5X PBS, 50 mM EDTA, 50 mM EGTA, and 1% carboxymethyl cellulose). The mixture was rotated overnight at 4°C. To decrease viscosity, cellulase was added to the samples for degrading carboxymethyl cellulose, followed by incubation at 37°C for 10 min. After incubation, the samples were washed once with PBS. The samples were processed for RNA extraction and western blotting.

### Western blotting

Samples were mixed with 4x Laemmli sample buffer (Bio-Rad) without a reducing agent and heated at 95°C for 5 min. Subsequently, the samples were loaded onto 4-20% gels (Bio-Rad), and electrophoresis was performed at 200V for 20 min. The proteins were then transferred onto polyvinylidene fluoride (PVDF) membranes using a wet transfer method. Membranes were blocked for 1 h at room temperature using Blocking One (Nacalai Tesque). Primary antibodies were added to the samples in Can Get Signal Solution 1 (Toyobo) and incubated overnight at 4°C. After washing three times with 1X PBST, the samples were incubated with secondary antibodies in Can Get Signal Solution 2 (Toyobo) for 1 h at room temperature and then washed three times in 1X PBST. Finally, imaging was performed using Amersham ECL Select (Cytiva) and ImageQuant LAS500 (Cytiva) software. The antibodies used in this study are listed in Supplementary Table S2 online.

### RNase protection assay

A spike-in control of the synthesized RNA (Pepper Mild Mottle Virus (*PMMoV*) partial sequence) was added to the crude sEVs sample at a final concentration of 5 pM. The sample was then treated with either 0.5 μg/μL RNase A (Nippon gene) or PBS for 20 min at 37°C. Total RNA was extracted using TRIzol LS Reagent (Thermo Fisher Scientific) following the manufacturer’s instructions. Reverse transcription qPCR (RT-qPCR) was performed using primers specific for *RAB13* and a spike-in sequence.

### Reverse transcription-quantitative PCR

Total RNA was extracted using TRIzol Reagent (Thermo Fisher Scientific) following the manufacturer’s instructions. The extracted total RNA was subjected to reverse transcription using ReverTraAce qPCR RT Mix/gDNA remover (Toyobo). RT-qPCR was performed using KOD SYBR qPCR Mix (Toyobo) on a QuantStudio 5 Real-Time PCR System (Thermo Fisher Scientific) following the manufacturer’s instructions. The abundance of the transcripts of interest (relative to *GAPDH*) and the EV/cell ratio were calculated. Primers used are listed in Supplementary Table S1 online.

### RNA sequencing and analysis

Total RNA was extracted from both cells and CD9 positive EVs using the TRIzol reagent (Thermo Fisher Scientific). Poly(A) RNA was isolated with the NEBNext Poly(A) mRNA Magnetic Isolation Module (New England Biolab) following the manufacturer’s protocol. The library was prepared using the GenNext RamDA-seq Single Cell Kit (Toyobo), according to the manufacturer’s instructions. Paired-end sequencing was performed with 150bp reads using NextSeq (Illumina). Raw reads were filtered using fastp (0.19.4)^15^ and mapped to the GRCh38.p12 reference genome using hisat2 (2.1.0)^16^. The FPKM was calculated using cufflinks (1.59.0)^17^. Genes enriched in sEVs that met the following criteria were extracted: FPKM in the EV sample > 100, log2 fold change (EV/cell) > 5, and p < 0.01.

### Plasmid construction and transfection

Complete or partial sequences of *RAB13* were chemically synthesized using an artificial gene synthesis service (GenScript) and inserted into either the pcDNA3.1+ or pcDNA3.1_N-eGFP vectors using the In-Fusion HD Cloning Kit (Takara). Following the manufacturer’s instructions, the Lipofectamine 3000 Transfection Reagent (Thermo Fisher Scientific) was used to transfect the plasmids into the cell lines. Subsequently, stably expressing cells were selected using Geneticin Selective Antibiotic (Thermo Fisher Scientific).

### Direct RNA sequencing and poly(A) tail length analysis

Total RNA was extracted from HEK293 cells expressing *eGFP::RAB13* (783-1164) sense or scrambled strands. Poly(A) RNA was isolated using a NEBNext Poly(A) mRNA Magnetic Isolation Module (New England Biolabs). Direct RNA sequencing libraries were prepared using a Direct RNA Sequencing Kit (Oxford Nanopore Technologies) and sequenced on MinION Mk1C with flow cells R9.4.1 (Oxford Nanopore Technologies). The sequenced data were base-called using guppy (5.1.12) with an HAC model. The resulting reads were mapped to the reference human transcript sequences from GRCh38.p12, to which the *eGFP* sequence was manually added, using minimap2 (2.24)^18^. Reads with a mapping quality of less than three were filtered out. The poly(A) tail lengths were calculated using tainfindr (1.3)^19^ in R (4.2.1)^20^. To analyze the relationship between poly(A) tail length and the EV/cell ratio, RNA-seq data obtained using a short-read sequencer were combined. The average poly(A) tail length of each gene was calculated and genes with zero FPKM values or zero poly(A) tail lengths in the cells were excluded from the analysis. Finally, the Pearson correlation coefficient between the poly(A) tail length and EV/cell ratio was calculated using R (4.2.1).

### Statistics

Statistical analyses were performed using R (4.2.1). All data were presented as mean ± standard deviation.

### Declaration of Generative AI and AI-assisted technologies in the writing process

During the preparation of this work, the authors used DeepL and ChatGPT to improve the readability and proofreading of the manuscript. After using these tools, the authors reviewed and edited the content as required and took full responsibility for the content of the publication.

## Acknowledgements

We would like to express our deep thanks to Dr. Kazuya Omi for his valuable review and suggestions for our manuscript. We also wish to acknowledge Ms. Marina Kishida for her help with the technical aspects of our work. In addition, we would like to thank Editage (www.editage.com) for English language editing.

## Author Contributions

Conceptualization: Y.O., K.T., Y.K.; investigation: Y.O., K.T., Y.K.; methodology: Y.O., K.T., Y.K.; formal analysis: Y.O.; visualization: Y.O., K.T.; writing-original draft: Y.O.; writing-review and editing: Y.O., K.T., Y.K.

## Data availability

Sequencing data are available from NCBI SRA with accession number PRJNA953174. All other data are available in the paper and supplementary information.

