## Supplementary Information for "A novel sorting signal for RNA packaging into small extracellular vesicles"

*Co-corresponding authors

### Supplementary Figures and Tables

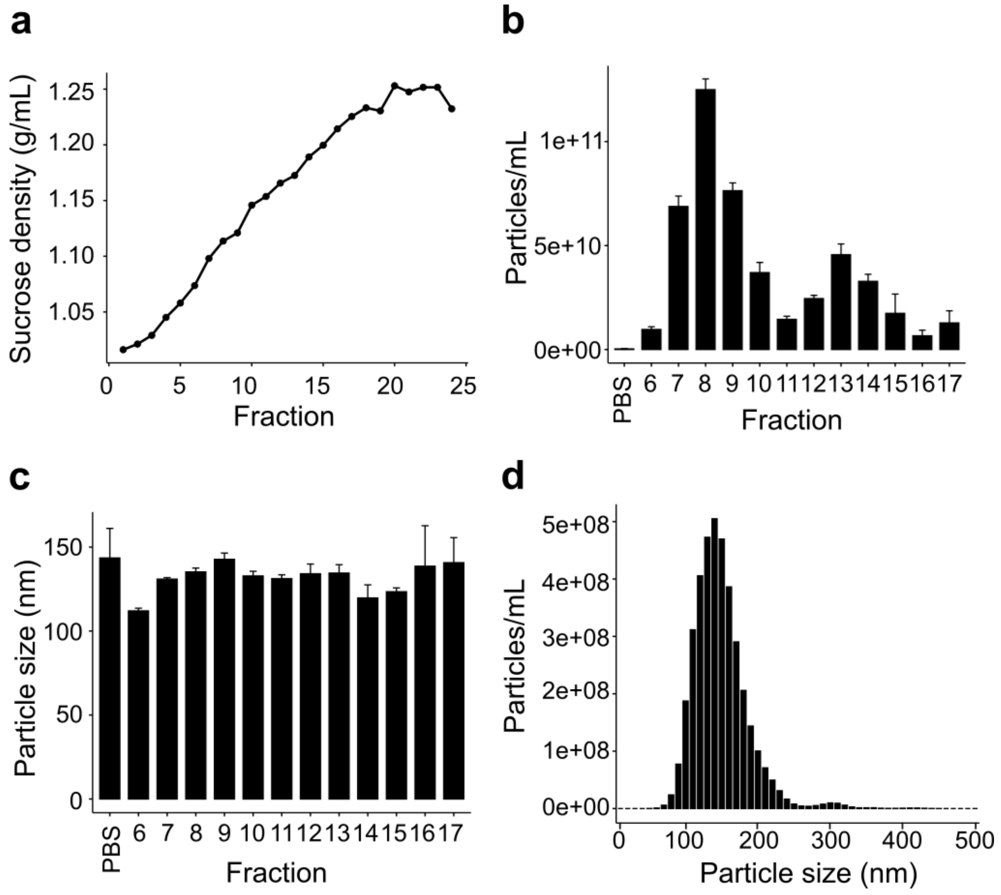


**Fig. S1 Fractions obtained through sucrose density gradient ultracentrifugation.** (a) Sucrose densities of each fraction. (b) Particle numbers in each fraction as measured by nanoparticle tracking analysis. Error bars are shown as ±standard error. (c) Average particle size in each fraction as measured by nanoparticle tracking analysis. Error bars are shown as ±standard error. (d) Histogram of particle size distribution of fraction 9.

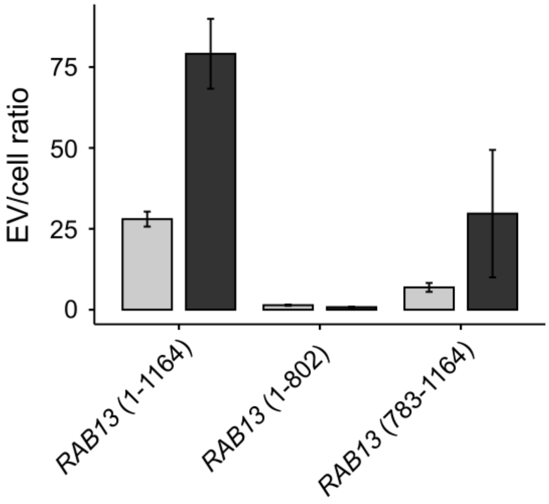


**Fig. S2 Enrichment level of *RAB13* complete or partial sequences in H1299 and MCF7 cell lines.** The transcripts of interest in CD9 positive EVs and cells were quantified and normalized to *GAPDH*, expressed as a ratio of transcript level in CD9 positive EVs to that in cells.

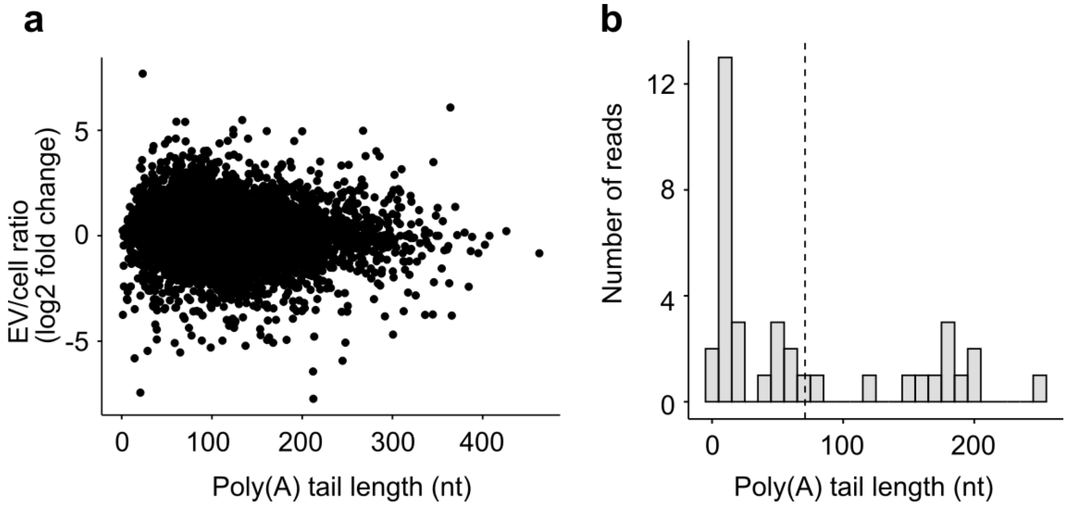


**Fig. S3 Association of EV enrichment level with poly(A) tail length from cells expressing the *eGFP* fusion gene with scrambled sequence of RAB13 (783-1164).** (a) Scatter plot showing the relationship between endogenous genes' poly(A) tail length and enrichment level in CD9 positive EVs (r = -0.11415). Poly(A) tail length data was obtained from cells expressing *eGFP::RAB13* (783-1164, scrambled sequence). (b) Poly(A) tail length distribution for the transgene extracted from cells expressing *eGFP::RAB13* (783-1164, scrambled sequence).

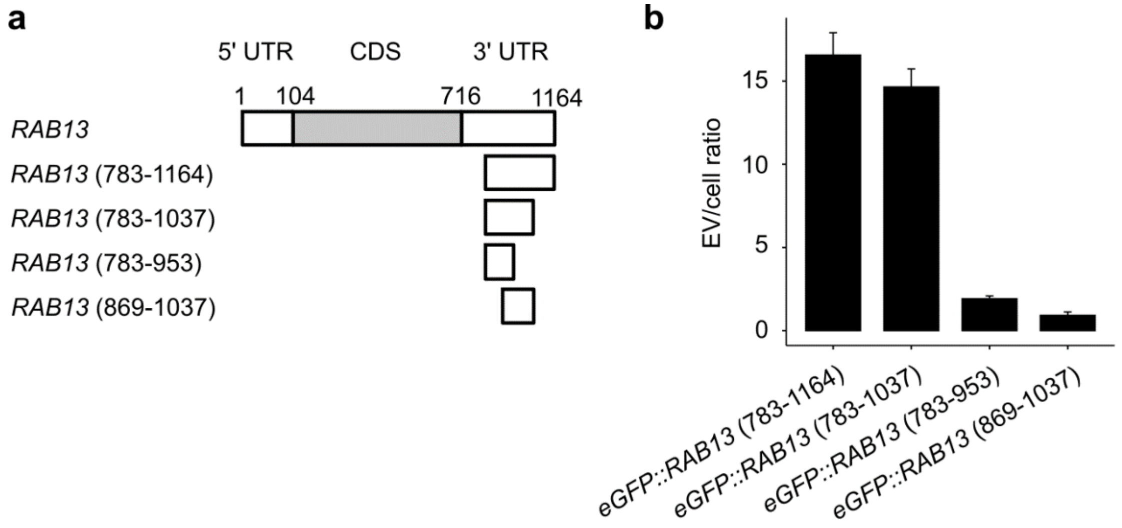


**Fig. S4. *RAB13* mRNA enrichment level of transgenes fused with *RAB13* 3'UTR subsequence in CD9 positive EVs.** (a) Diagram of the *RAB13* mRNA and its partial sequences. The indicated number in parentheses corresponds to the sequence positions of *RAB13*. The black bar in the 3' UTR shows polyadenylation signal. (b) Enrichment level of transgenes in the CD9 positive EVs from cells expressing the partial sequences of *RAB13* fused with *eGFP*.

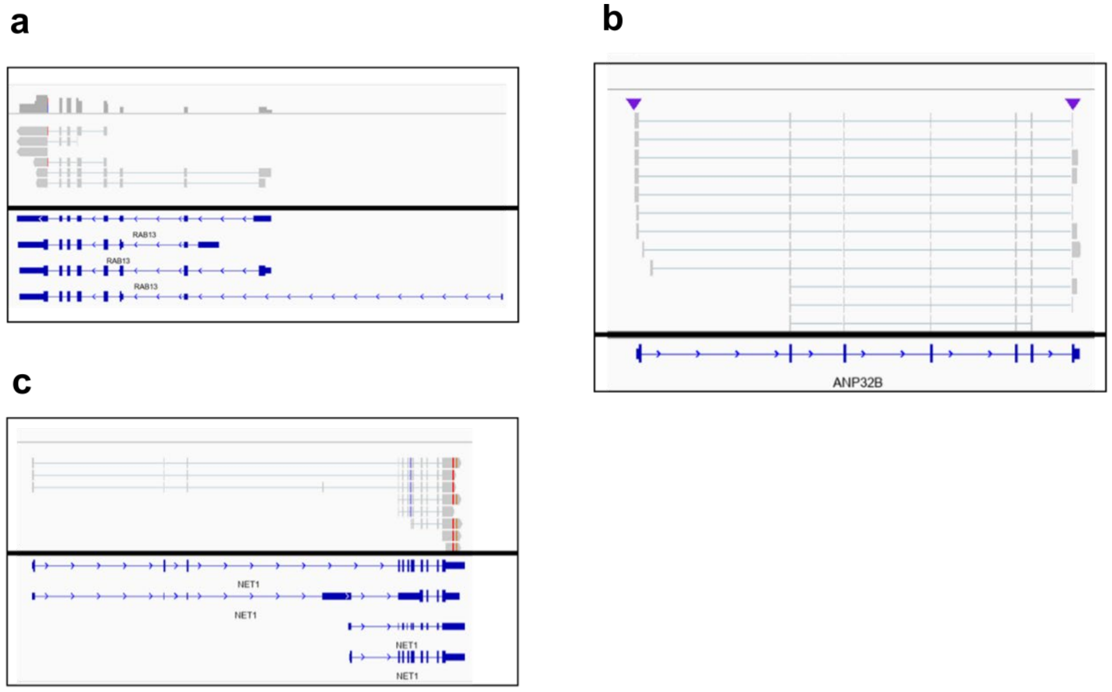


**Fig. S5. Full-length reads of enriched transcripts detected in CD9 positive EVs.** Mapped reads to (a) *RAB13*, (b) *ANP32B*, and (c) *NET1* reference sequences were visualized using Integrated Genome Viewer.

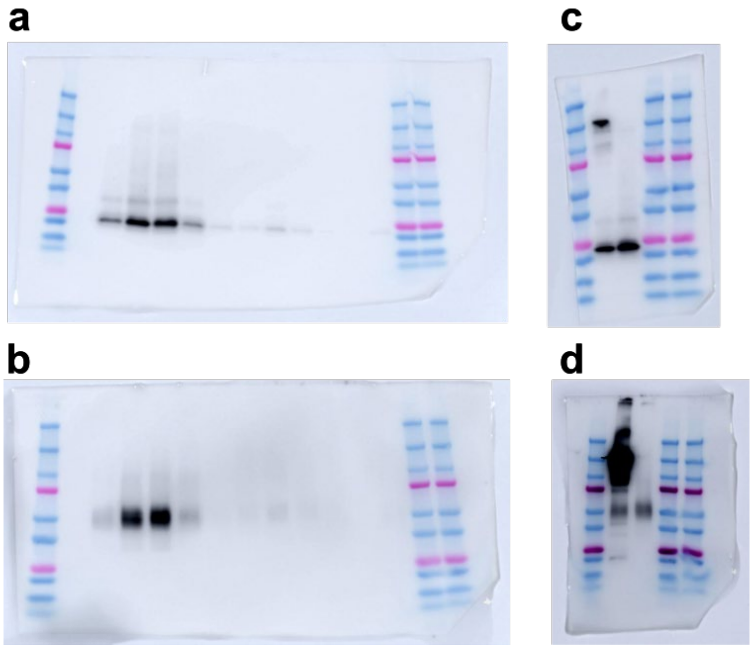


**Fig. S6. Raw western blot images.** (a) CD9 and (b) CD63 in Fig. 1a. (c) CD9 and (d) CD63 in Fig. 1b. The upper bands of (c) and (d) correspond to anti-CD9 antibody conjugated magnetic beads.

#### Table S1. Primers and sequences

| Primers and sequences | Sequence (5' → 3') | Description |
|---|---|---|
| *RAB13* Fw<br>*RAB13* Rv | AGACAATAACTACTGCCTACTACCGTG<br>GAGCACTTGTTGGTGTTCTTCTTGTCA | For quantifying endogenous *RAB13* (Fig. 1d-f) |
| *ANP32B* Fw<br>*ANP32B* Rv | GTCAGTGAGGAGGAAGAAGAATTTGG<br>GTAGATTACCAAGAGGGACTACATGG | For quantifying endogenous *ANP32B* (Fig. 1d) |
| *NET1* Fw<br>*NET1* Rv | CAGTCCAAGAGCTAGTCCTAGAAGAG<br>CTTCTACCTGAGTAACACTGGAAACTG | For quantifying endogenous *NET1* (Fig. 1d) |
| *LUC7L3* Fw<br>*LUC7L3* Rv | CAGAGAACAAGATAGAAAATCCAAGG<br>AACAGTAACAAAAAGCCCTAATCAAA | For quantifying endogenous *LUC7L3* (Fig. 1d) |
| *ACTG1* Fw<br>*ACTG1* Rv | CTTCCTTCCTGGGTATGGAATCTTG<br>CAAATTTCTATTCTCAATTAACCCATG | For quantifying endogenous *ACTG1* (Fig. 1d) |
| *PMMoV* Fw<br>*PMMoV* Rv | GAGTGGTTTGACCTTAACGTTTGA<br>TTGTCGGTTGCAATGCAAGT | For quantifying spike-in RNA (Fig. 1f) |
| pcDNA3.1 Fw<br>pcDNA3.1 Rv | GAGCTCTCTGGCTAACTAGAGAACCC<br>CAGACAATGCGATGCAATTTCCTC | For quantifying partial sequences of *RAB13* (Fig. 2b, S2) |
| *GAPDH* Fw<br>*GAPDH* Rv | ACTTTGTCAAGCTCATTTCCTGGTATGAC<br>GGTACTTTATTGATGGTACATGACAAGGTG | For quantifying *GAPDH* as internal control (Fig. 1,2, S2, S4) |
| *eGFP* Fw<br>*eGFP* Rv | GACTGGGTGCTCAGGTAGTG<br>CAAGATCCGCCACAACATCG | For quantifying fusion genes (Fig. 2c, S4) |
| *PMMoV* spike-in sequence | GCAGCAAAGGUAAUGGUAGCUGUGGUUUCAAAUGA<br>GAGUGGUUUGACCUUAACGUUUGAGAGGCCUACCG<br>AAGCAAAUGUCGCACUUGCAUUGCAACCGACAAUU<br>ACAUCAAAGGAGGAA | For Spike-in control (Fig. 1f) |

#### Table S2. Antibodies

| Antibodies | Usage | Source | IDs |
|---|---|---|---|
| Mouse monoclonal anti-CD9 | Immunoaffinity purification of sEVs | In-house | - |
| Mouse monoclonal anti-CD9 | Western blot | CosmoBio | SHI-EXO-M01 |
| Mouse monoclonal anti-CD63 | Western blot | In-house | - |
| Rabbit Anti-Mouse Immunoglobulins/HRP | Western blot | Dako | P0260 |

### Supplementary Methods

#### Nanoparticle tracking analysis

Fractions obtained through sucrose density gradient ultracentrifugation and fractionation were diluted at 20-fold or 40-fold and subjected to nanoparticle tracking analysis using a NanoSight LM10 (Malvern Panalytical). The results were evaluated using Nanoparticle Tracking Analysis software (3.1).

#### Full length RNA sequencing and analysis

The conditioned medium from HEK293 cells was centrifuged at 2,000 x g for 5 minutes at 4 °C, followed by filtration through a 0.45 μm filter. The samples were enriched using an AMICON ULTRA-15 100 KDa cutoff (Merck Millipore) and resuspended in PBS. Subsequently, the enriched samples were cautiously layered onto 30w/v% sucrose solution present in ultracentrifugation tubes and subjected to ultracentrifugation at 100,000 x g for 3 hours at 4 °C. The collected sucrose solutions were mixed with PBS and underwent ultracentrifugation at 100,000 x g for 1 hour for washing purposes. Total RNA was extracted using TRIzol LS Reagent (Thermo Fisher Scientific) following the manufacturer's instructions.

Library preparation and sequencing were performed according to Procedure-Checklist-Iso-Seq-Express-Template-Preparation-for-Sequel-and-Sequel-II-Systems version02 (PacBio). Briefly, cDNA synthesis and amplification were carried out using the NEBNext Single Cell/Low Input cDNA Synthesis & Amplification Module (NEB), with primers from the Iso-

Seq Express Oligo Kit (PacBio). The PCR cycle consisted of 20 cycles. The amplified cDNA was size-selected to remove fragments less than 1 kb using the ProNex® Size-Selective Purification System (Promega). Libraries were prepared with the SMRTbell Template Prep Kit 2.0 (PacBio) and sequenced on the PacBio Sequel I platform. Analysis was performed using the Iso-Seq Analysis in SMRT Link. Briefly, circular consensus sequences (CCS) reads were generated from the sequenced reads and primer sequences, and the poly(A) tail and concatemers were removed. The reads were then clustered, and a consensus sequence was generated for each read cluster. The polished reads were mapped to a reference sequence and visualized using the Integrated Genome Viewer.
